# The Lion’s Share: Implications of carnivore diet on endangered herbivores in Tsavo

**DOI:** 10.1101/2023.07.08.548222

**Authors:** Eliza King, Stotra Chakrabarti, Fredrick Lala, Stephen Nyagah, Grace Waiguchu, Patrick I. Chiyo, Joseph Kimaile, Richard Moller, Patrick Omondi, Aaron Morris, Joseph K. Bump

## Abstract

Predation by mammalian carnivores can have cascading, regulatory effects across ecological communities. An understanding of predator diet can therefore provide crucial information regarding their ecology and conservation, as well as their impacts on prey populations. Using scats collected between 2019 and 2023, coupled with estimates of prey abundance from aerial surveys, we characterized lion prey-consumption and preference in Tsavo, Kenya. A lion-specific biomass model applied to prey frequencies in scats revealed that more than 85% of lion diet consisted of large ungulates weighing over 150 kg. While large ungulates were also preferred prey items in terms of their availability, we found a disproportionately high consumption and preference of lions for the endangered hirola and Grevy’s zebra— species that were introduced in Tsavo as part of ex-situ conservation programs. Hirola and Grevy’s zebra populations have historically remained small in Tsavo despite strong recovery efforts, and our results likely indicate a disproportionate impact of lion predation on these small but crucial populations. Preferential predation, coupled with high availability of alternative prey, may trap hirola and Grevy’s zebra within a *predator-pit*. Our findings have strong implications for understanding lion diet, optimal foraging, and the potential effects predators have on endangered prey species in a landscape of critical conservation importance.

## II. Main Text

### Introduction

Carnivores are at the forefront of conservation efforts because of their threatened status, yet they play crucial umbrella roles within their ecosystems (Gittleman et al., 2001; Dalerum et al., 2008, Ripple et al., 2014). Carnivore charisma and connection with the human psyche often make them flagship species in conservation (Gittleman et al., 2001; Ducarme et al., 2013). The tiger (*Panthera tigris*), lion (*Panthera leo*), gray wolf (*Canis lupus*), and cheetah (*Acinonyx jubatus*) have been the face of projects focused on habitat conservation, ecosystem recovery, and conservation introductions (Bangs & Fritts, 1996; Jhala et al., 2019; Packer, 2019; Jhala et al., 2021). While such conservation policies are currently aimed towards the restoration of carnivore populations through protection and augmentation, their recovery can lead to increased conflicts with humans (e.g. Asiatic lions in India, Jhala et al., 2019; brown bears, *Ursus arctos*, in the Pyrenees mountains, Piedallu et al., 2016) and reduction of endangered prey species, especially when alternative prey is abundant (e.g. bighorn sheep *Ovis canadensis* predation by puma *Puma concolor* when sympatric mule deer *Odoceilus hemionus* is abundant, Johnson et al., 2012). Because of such layered effects mediated by carnivores, a critical understanding of their ecology and behavior remains key for the effective management of ecosystems (Boitani & Powell, 2012).

Lions are iconic for their connection to human culture and their apex position in ecosystems, making them mascots of conservation wherever they occur (Roemer et al., 2009). However, lions are also one of the top species in conflict, threatening human lives and livelihood (Ray et al., 2005). Direct persecution and detrimental habitat alteration has limited the range of extant African lions to sub-Saharan Africa, although historically their range included almost the entirety of the continent (Ray et al., 2005). Their numbers have plummeted by more than 75% in the past 50 years, with a range contraction over 90% (Ripple et al., 2014; Loveridge et al., 2022). Tsavo Conservation Area (hereafter TCA), located in southeastern Kenya, is a semi-arid, drought-prone landscape that remains one of the lion strong-holds in East Africa (Henschel et al., 2020). This landscape also forms a crucial connection between two major African biomes— mesic grassland savannah to the south and semiarid bush savannah to the north (Henschel et al., 2020). About 700 lions live in the contiguous landscape of Tsavo (covering both East and West Tsavo National Parks), yet lion ecology has been traditionally understudied in this landscape (barring mane growth, Kays & Patterson, 2002, and human-eating tendencies of 2 infamous male lions, DeSantis & Patterson, 2017). This contrasts with other well-studied East African populations such as in Serengeti and Ngorongoro (reviewed in Packer et al., 2019).

TCA is also home to one of the last remaining, albeit crucial, populations of the critically endangered hirola (*Beatragus hunteri*). The hirola population is restricted to a small area of their natural range at the Kenyan-Somali border, but were introduced to Tsavo East National Park in 1962 and 1996 as insurance against imminent extinction triggered by massive population declines (reviewed in Probert et al., 2014). The hirola population in Tsavo has remained small, and such herbivore populations can be prone to the detrimental effects of opportunistic predation when alternate prey is abundant (Johnson et al., 2012). Similar to the hirola, the endangered Grevy’s zebra (*Equus grevyi*) was also introduced to TCA in 1964 and 1977 with similar conservation interests, but the population has remained small (Githiru, 2017). Multiple factors can impede recovery of introduced herbivores, and among them, predation features as a plausible cause for these two crucial populations in Tsavo (Evans, 2011 and Probert et al., 2014).

To understand the general ecology of lions in this important population, as well as their potential predation on endangered hirola and Grevy’s zebra, we apply scat analysis using biomass models coupled with prey abundance estimates to present information on lion prey consumption and preference in Tsavo East National Park, an area where these two herbivore populations are currently restricted. Our results have important bearings on predation ecology on endangered herbivores in a conservation-significant landscape.

### Study Area

We conducted the study in the TCA, 1°59′S-4°8′S 37°45′E-39°16′E, which spans over 42,000 km^2^ and includes Tsavo East and West National Parks, Chyulu National Park, South Kitui National Reserve, Taita, Galana and Kulalu Ranches, Mkomazi National Park, and privately owned ranches/reserves (Lala et al., 2021). Tsavo East comprises an area of 13,747 km^2^. It’s a typically semiarid landscape with a dry season spanning from January through early March and a cooler season from June to October. Annual temperature ranges from 20°C to 30°C, with an annual precipitation ranging between 200 to 700 mm (Lala et al., 2021).

A few major rivers run through the area including the Galana, Tsavo, and Voi. Much of the landscape is open and shrubbed grasslands, while a mosaic of riparian habitats predominates along the rivers. Typical wild herbivores include elephants (*Loxodonta africana*), giraffe (*Giraffa camelopardis*), plains and Grevy’s zebra (*Equus* spp.), Cape buffalo (*Syncerus caffer*), and a reintroduced population of hirola, while carnivores such as lions, cheetahs, leopards (*P. pardus*), African wild dogs (*Lyacon pictus*), and spotted hyenas (*Crocuta crocuta*) are common. A list of the mammalian and avian species richness and diversity of TCA can be found in Lack et al. (1980), Lepage (2004), and Tóth et al. (2014). Tsavo is a keystone biodiversity habitat whose important connection with Amboseli National Park creates the larger Tsavo-Amboseli ecosystem.

## Methods

### Scat Collection and Processing

We opportunistically collected 74 lion scats from Tsavo East National Park between November 2019 and February 2023 (**Figure 1**). All scats were geo-referenced in the field. To avoid misidentification of scats, we adopted one or a combination of the following field techniques: scats were collected either i) when a lion was observed defecating, ii) from known lion-used trails after checking for additional evidence such as the presence of tracks and scrapes, and iii) scats that were >35 mm in diameter to reduce confusion with leopard scats. Further, we relied on the ecological knowledge of experienced field personnel from the Wildlife Research and Training Institute and Kenya Wildlife Service rangers who are familiar with the area. We did not include ambiguous or confusing samples. This constitutes a conservative approach that increases the certainty that all of the samples used for this analysis were likely lion scats.

**Figure 1.**
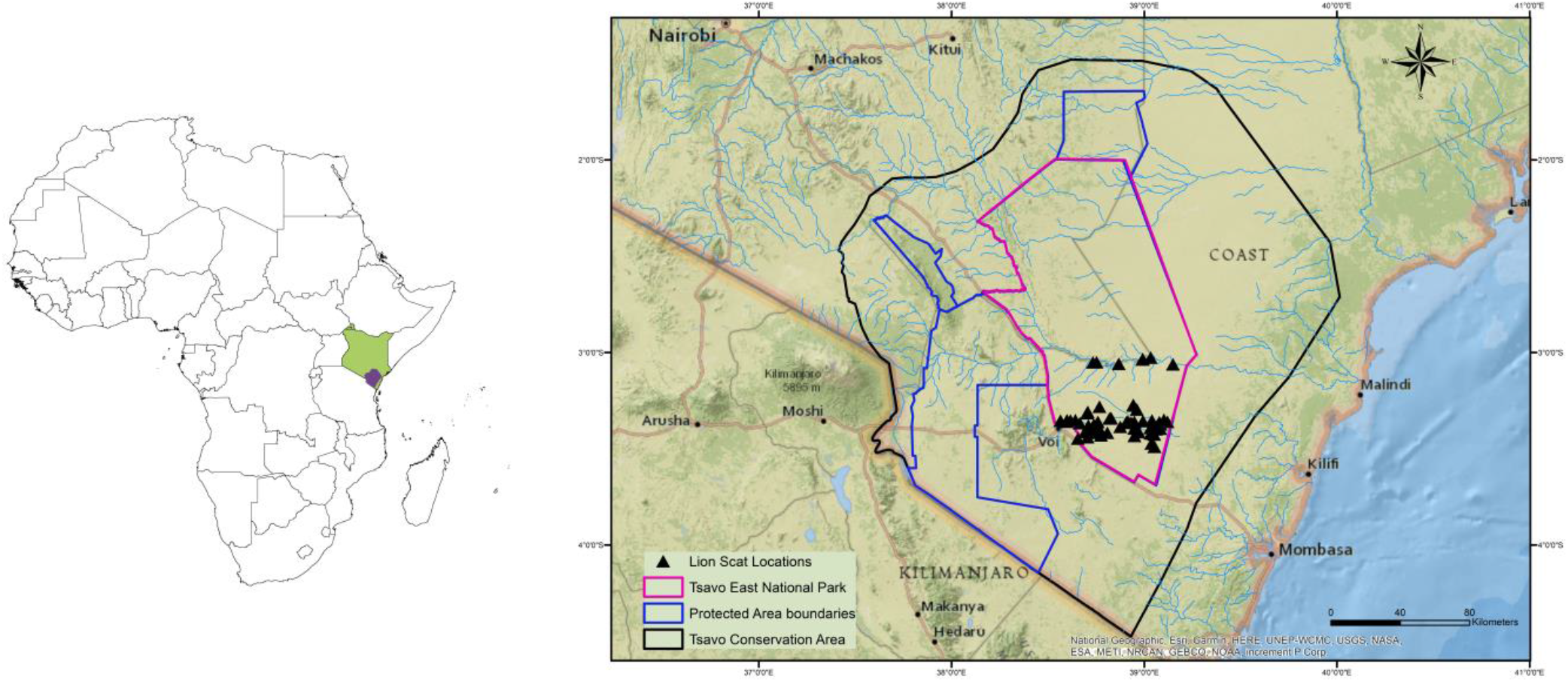
A map showing the relative positions of Kenya (filled green, Left) and Tsavo Conservation Area (filled purple, Left) in Africa. The magnified map in the inset (Right) points to the locations of the 74 lion scat samples that we collected and analyzed from Tsavo East National Park between 2019 and 2023.

Each collected scat was then sun dried before being washed to collect the prey remains in a sieve. We randomly selected 20 prey hairs from each washed sample, and each hair was placed on a slide. Prey hairs were then identified to the species level under a light microscope using a reference key for mammalian hair from the area as described by Perrin & Campbell, 1979. The number of prey hairs belonging to a specific species relative to the complete 20-hair set from each scat represented the per whole scat equivalence (WSE) of that prey. The cumulative per-scat equivalence of a prey species across all 74 scats was used to compute biomass consumption.

### Prey Consumption

We calculated the total biomass consumption of each prey species found in the scats through a lion specific biomass model developed by Chakrabarti et al. (2016): y = 4.105 − 3.116*exp*^−0.032*x*^, where *x* is the average prey weight and *y* is the prey biomass consumed per scat. The biomass consumed per scat (*y*) for each prey species was then multiplied with the total scat equivalence of that species in our collected samples, resulting in the estimated actual biomass consumed of that particular prey in our collected samples. For prey weights, we used 3/4 of the mean adult female body weight following Schaller (1972) and Hayward & Kerley (2005) because it accounts for the weights of juveniles and sub-adults in the population. This means it is a better representation of the average weight of a prey population available to a predator. Prey weights were adopted from Stewart (1963), Sachs (1967), Myers et al. (2008), Pinto (2018), and Hayward & Kerley (2005). We further calculated the relative biomass consumed for each prey as a proportion of total consumption to understand the importance of each prey species in lion diet.

### Prey Selection

We used recent herbivore aerial census data for the TCA (Waweru et al., 2021) to calculate Tsavo East-specific abundance estimates for prey species occurring in the lion scats. We then calculated the relative abundance of each prey species as an index of availability, where the abundance of each prey was converted into a proportion of the total abundance of all the species occurring in the lion scats in Tsavo East. We could not include dikdik (*Madoqua* sp.) in this prey selection analysis owing to the lack of a population estimate. Their small size, solitary life-history, and concealment under dense habitat precluded accurate aerial counting. However, dikdik are not usually included in lion diet (Hayward & Kerley, 2005; current study), and thus we are confident that such an exclusion has not significantly altered prey selection indices for lions in our study. We also excluded elephant from this part of our analysis as they only appeared in one scat sample and did not represent a significant part of lion diet. From the relative biomass consumed and relative prey density, we computed Jacobs’ selectivity index for each prey species occurring in lion scats using the *dietR* package in R (V.4.2.1). Jacob’s selectivity indices (D) range from +1 to -1, indicating maximum preference and maximum avoidance respectively, while a value closer to 0 represents random selection of a prey item (Jacobs, 1974).

## Results

### Scat collection and processing

Based on the cumulative species occurrence plot, we found our sampling to be adequate to characterize lion diet in Tsavo East because new species occurrence in scats satiated at ∼20 scats (**Figure S1**). We identified 16 prey species from the scats; elephant being the largest and dikdik being the smallest on the size spectrum (**Table 1**).

**Table 1.** Lion prey consumption and preference in Tsavo East National Park. Table with body weights, whole scat equivalence (WSE), biomass consumed per scat (Y), total and relative biomass consumed, total and relative abundance, and Jacobs’ index values (D) for 16 prey species found in 74 lion scats collected between 2019 and 2023. Species in bold represent significantly preferred prey items, with a threshold of preference/avoidance set at +/- 0.3. Prey weight corresponds to 3/4 of mean adult female weight. Dikdik could not be included in the prey preference analysis owing to the lack of a population estimate, and elephant was excluded as elephant remains were found only in one scat sample.

| Species | Weight (kg) | WSE | Y<br>(kg/scat) | Biomass Consumed<br>(kg) | Rel. Consumption<br>(%) | Abundance | Rel. Abundance<br>(%) | D |
| --- | --- | --- | --- | --- | --- | --- | --- | --- |
| **Buffalo** | 432 | 23.10 | 4.10 | 94.83 | 33.47 | 1351 | 10.85 | **0.51** |
| **Waterbuck** | 188 | 12.10 | 4.10 | 49.58 | 17.50 | 83 | 0.67 | **0.93** |
| Giraffe | 550 | 9.40 | 4.10 | 38.59 | 13.62 | 998 | 8.01 | 0.26 |
| Plains zebra | 175 | 8.80 | 4.09 | 36.02 | 12.71 | 3580 | 28.75 | -0.38 |
| **Warthog** | 45 | 4.40 | 3.37 | 14.81 | 5.23 | 136 | 1.09 | **0.66** |
| Eland | 345 | 3.25 | 4.10 | 13.34 | 4.71 | 668 | 5.36 | -0.06 |
| **Grevy's zebra** | 190 | 1.90 | 4.10 | 7.79 | 2.75 | 14 | 0.11 | **0.92** |
| **Hirola** | 90 | 1.70 | 3.93 | 6.68 | 2.36 | 68 | 0.55 | **0.63** |
| Impala | 30 | 1.90 | 2.91 | 5.53 | 1.95 | 837 | 6.72 | -0.55 |
| Hartebeest | 95 | 1.15 | 3.96 | 4.55 | 1.61 | 1174 | 9.43 | -0.71 |
| Beisa oryx | 158 | 1.05 | 4.09 | 4.29 | 1.51 | 1930 | 15.50 | -0.82 |
| Gerenuk | 44 | 1.25 | 3.34 | 4.18 | 1.47 | 132 | 1.06 | 0.17 |
| Grant's gazelle | 38 | 0.30 | 3.18 | 0.95 | 0.34 | 1352 | 10.86 | -0.94 |
| Lesser kudu | 99 | 0.15 | 3.97 | 0.60 | 0.21 | 131 | 1.05 | -0.67 |
| Dikdik | 4 | 0.25 | 1.36 | 0.34 | 0.12 |  |  |  |
| Elephant | 1600 | 0.30 | 4.11 | 1.23 | 0.43 |  |  |  |

### Prey Consumption

The majority of lion diet was composed of herbivores that weighed over 150 kg, with zebra, giraffe, buffalo, and waterbuck accounting for more than 70% of biomass intake (**Figure 2**). While small antelopes, such as gerenuk and Grant’s gazelle were rarely consumed (1% of the total biomass intake), warthog comprised ∼5% of lion diet. More than 5% of their diet consisted of the endangered Grevy’s zebra and hirola, species that are rare in the landscape and are of critical conservation concern.

**Figure 2.**
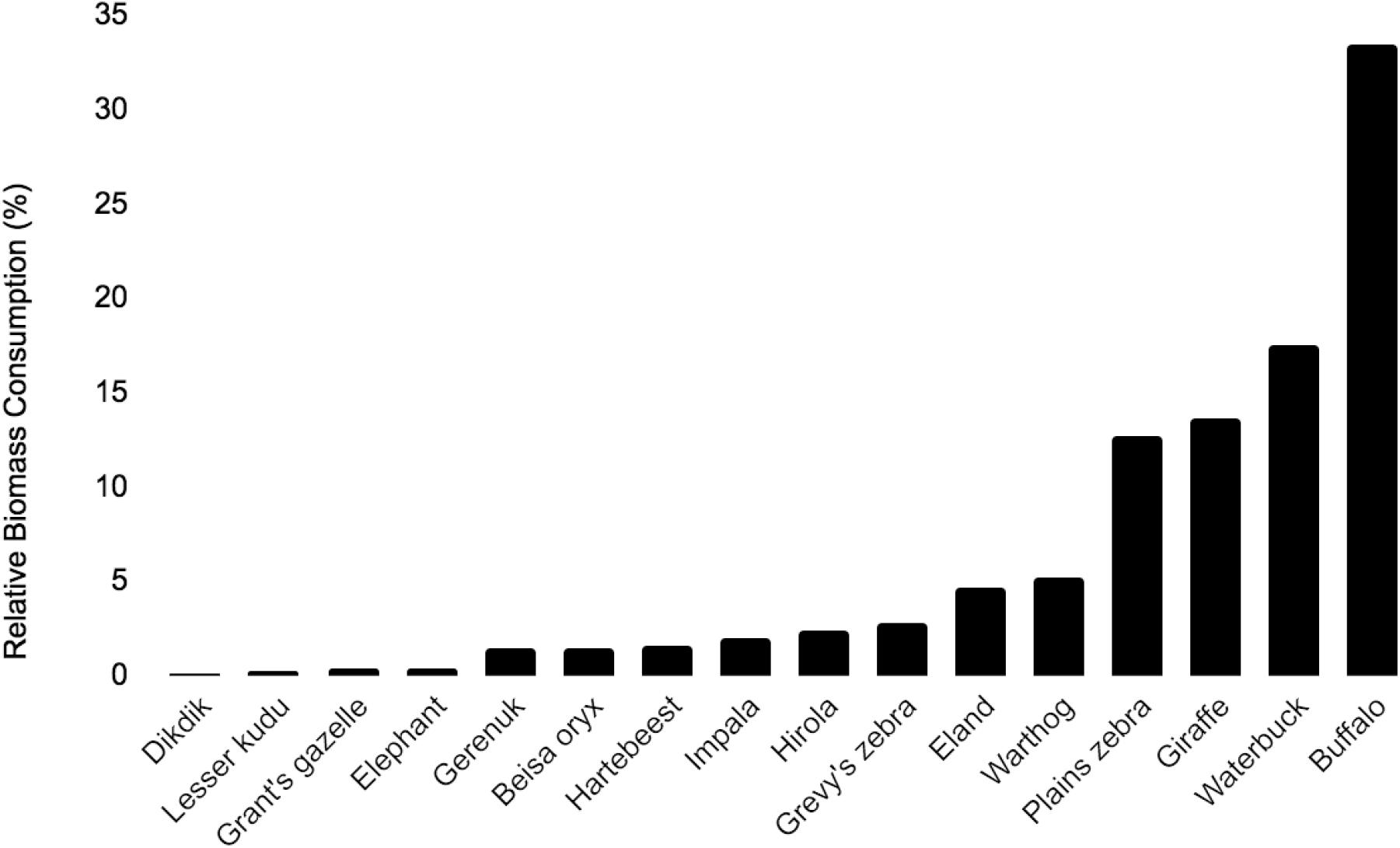
Relative biomass consumption of 16 prey species occurring in 74 lion scat samples collected from Tsavo East National Park, Kenya between 2019 and 2023. Each bar represents the contribution of a particular prey item relative to the total consumption, expressed as a percentage. Dikdik was consumed the least while buffalo was consumed the most. Biomass consumption of prey species was calculated using a lion-specific biomass model developed by Chakrabarti et al. (2016).

### Prey Selection (Jacob’s Index)

Based on prey availability, waterbuck, Grevy’s zebra, warthog, hirola, and buffalo were strongly preferred by lions, while Grant’s gazelle, Beisa oryx, hartebeest, impala, and kudu were avoided. Plains zebra, eland, giraffe, and gerenuk were consumed randomly/as per their availability (**Figure 3**).

**Figure 3.**
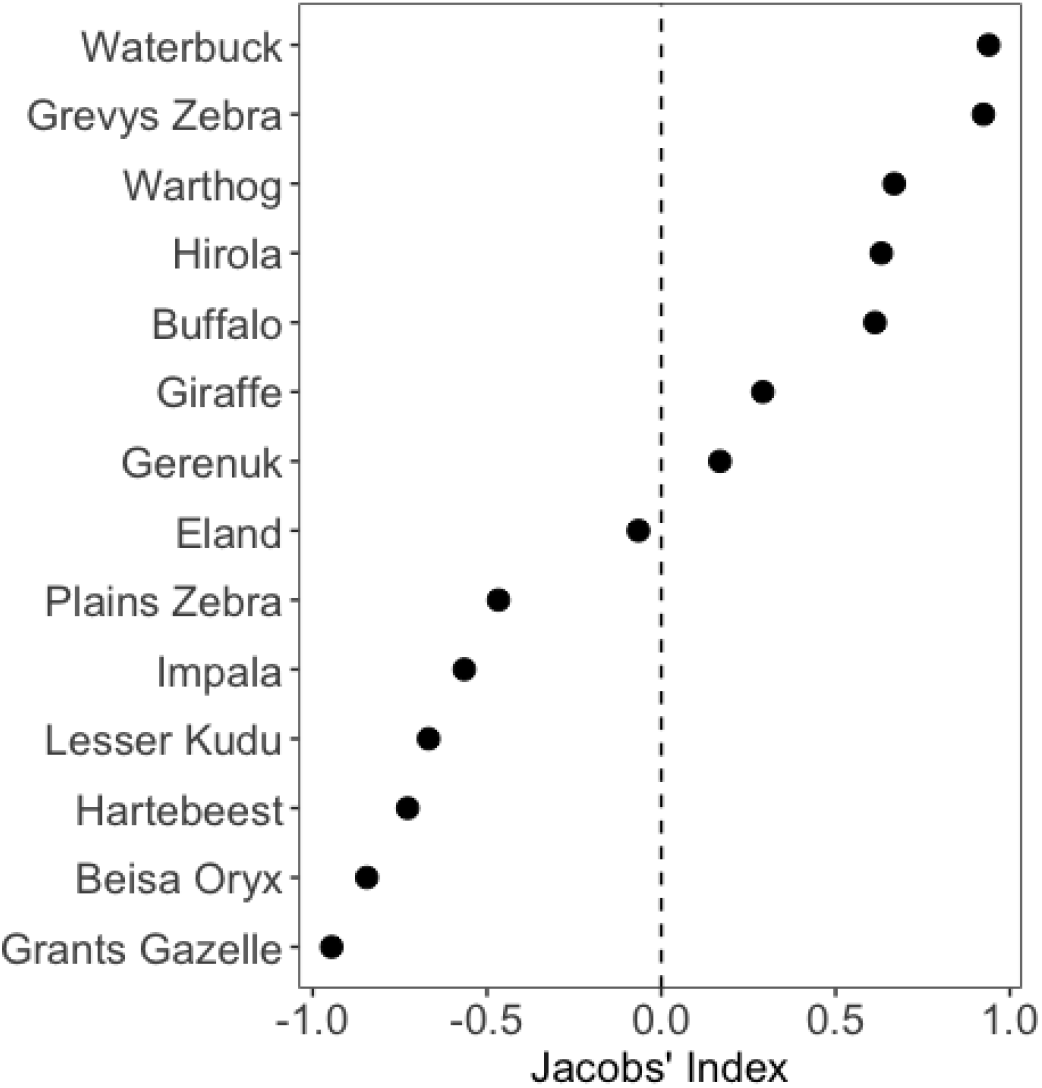
Jacobs’ selectivity indices (D) for prey species (excluding dikdik and elephant) occurring in 74 lion scats collected opportunistically from Tsavo East National Park, Kenya 2019-2023. Values <-0.3 indicate an avoidance and values >0.3 suggest a preference for the corresponding species, while values in between +/- 0.3 indicate that the corresponding species were chosen randomly/in proportion to their availability. Waterbuck, Grevy’s zebra, warthog, hirola, and buffalo were among the most preferred prey items.

## Discussion

A comprehensive characterization of an animals’ diet is critical for understanding species ecology. This is even more crucial for an apex carnivore because their predation effects can have cascading implications for the entire ecosystem. We investigated diet and prey selection of lions in Tsavo East National Park, Kenya using scat analysis through a lion-specific *biomass conversion model* and estimation of prey abundance. All earlier studies on African lion diet have either relied on direct observations and/or frequency of occurrence of prey remains in scats (reviewed in Hayward & Kerley, 2005), which have their limitations of either overestimating large or small prey respectively (Chakrabarti et al., 2016). *Biomass conversion models* account for differential prey digestibility, and hence provide a more accurate picture of carnivore diet from scats (Ackerman et al., 1984; Chakrabarti et al., 2016).

Our results reveal that Tsavo lions include a wide size range of mammalian prey species in their diet (from dikdiks to elephants), however they typically rely on large ungulates, with over 85% of their diet consisting of herbivores that weigh more than 150 kg (**Figure 5**). Our results also reflect similar trends in lion diet across their global range (Hayward & Kerley, 2005), as well as from the adjacent population in Amboseli (Courtois, 2015). Lion prey selection in Tsavo is in consonance with optimal foraging theory and biomass scaling that maintains that predators >21 kg should ideally consume large prey to maximize energetic returns (Krebs & Davies 2009; Carbone et al., 1999). However, the preferred prey weight range of Tsavo lions is typically on the heavier side, with multiple large- and mega-herbivores included among their prey (**Figures 3 & 4**). This is probably because lions are social carnivores, and their preference for large bodied ungulates results from the need of maximizing optimal foraging benefits for their group as a whole. However, lions rarely hunt cooperatively, with individual lions capable of successfully procuring most prey sizes (Packer, 1986; Packer et al., 1990). Cooperative and coordinated hunting in lions has been documented almost exclusively from Etosha National Park in Namibia, where lions hunt large and fleet-footed herbivores such as giraffes, and the hunting success of single lions is very low (Stander, 1992). Our analysis suggests that lions in Tsavo frequently consume formidable prey species such as buffalo and giraffe, warranting direct observation-based investigations of possible cooperative hunting strategies in this lion population. Furthermore, carnivores are known to disproportionately select juvenile and sub-adults of prey species that are dangerous as adults, and such direct observations can shed more light into age and sex-specific predation rates. Tsavo lions are also known to scavenge on elephant carcasses, which become available because of droughts and poaching (Morris et al., 2023). Surprisingly we found elephant remains in only one scat, which might be attributable to our study period coinciding with no major droughts coupled with disproportionate consumption of flesh and organs from large carcasses by predators; the latter being devoid of hair do not appear in scats (Chakrabarti et al., 2016). Going forward, the current analysis coupled with DNA-metabarcoding can potentially add crucial value to species identification (Thuo et al., 2019).

**Figure 4.**
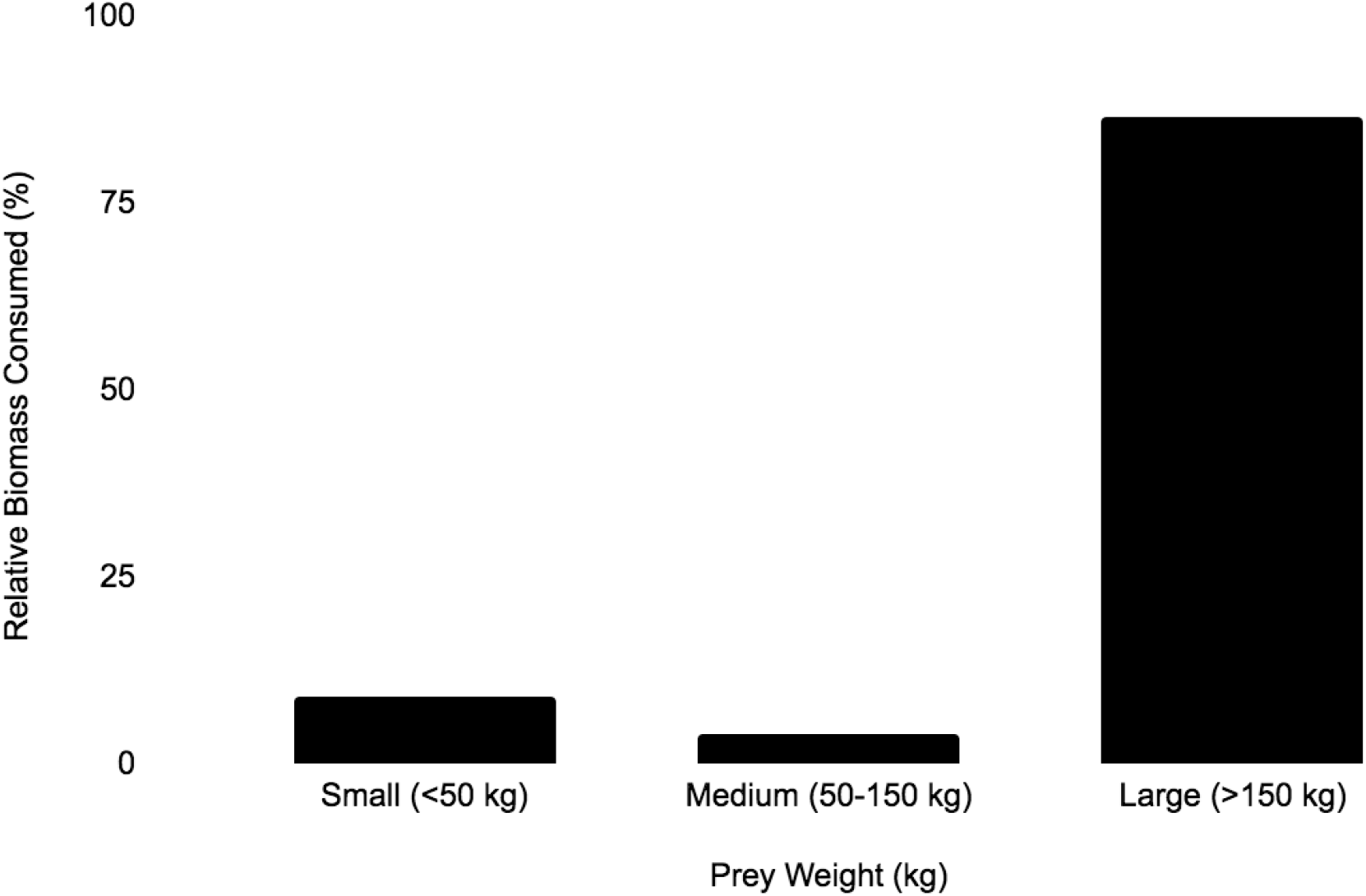
Relative biomass consumption of lion prey segregated into weight categories (small, medium, and large) in Tsavo East National Park, Kenya 2019-2023. Large prey constitutes the majority (>85%) of lion diet. Prey weight corresponds to ¾ mean adult female weight.

While large and abundant ungulates such as buffalo and waterbuck were among the most consumed and preferred prey items for Tsavo lions, we also found considerable consumption of hirola and Grevy’s zebra – both species are rare and restricted in spatial-pockets in the landscape. Cumulatively these species constituted almost 5% of lion diet and were highly preferred. The introduced hirola and Grevy’s zebra populations in Tsavo are of critical conservation importance due to relatively low numbers of the species in its natural range - the Tsavo hirola population constitutes nearly 20% of its global abundance (Probert et al., 2014; Waweru et al., 2021). The hirola and the Grevy’s zebra populations in Tsavo has remained small, and their recovery is a management priority. While hirola and Grevy’s zebra conservation in Tsavo has been at the forefront for decades, little is known about the potential stressors that have historically kept these population at low numbers (Githiru, 2017; Jowers et al., 2020). Another study, which reported frequency of occurrence of prey remains in lion scats also suggest noteworthy predation of hirola by lions in Tsavo (Evans 2011). Any small prey population can be disproportionately vulnerable to predation when alternative prey is available (Sinclair et al., 1998; Johnson et al., 2012). In the Tsavo system, large ungulates are fairly abundant and available for lions to subsist upon, as indicated in our current study. Such abundant alternative prey can trigger considerable *opportunistic* predation on hirola and Grevy’s zebra by predators/lions that are subsidized by other prey. Continued predation in magnitudes that are detrimental to population recruitment, can potentially trap these two species in a low population equilibrium or a ‘predator pit’ (Messier, 1994; Sinclair et al., 1998). Based on our analysis that shows lions in Tsavo East significantly predate on hirola and Grevy’s zebra, we recommend telemetry based observational studies on the large carnivore guild in TCA along with population monitoring of hirola and Grevy’s to understand age- and sex-specific predation, as well as recruitment. Such a study will be crucial to outline strategies for the effective management of these endangered herbivores in the Tsavo ecosystem.

## Author contributions

Study conception and design: SC, FL, JKB, AM; Field & lab work: SN, GW, PIC, JK, FL; Data analysis: EK, SC; Writing: EK, SC with inputs from JKB, FL, PO & RM; Funding: JKB, RM, SC; Review and approval: all authors; Project supervision: SC, JKB, FL, PO

## Acknowledgements

We would like to thank the Wildlife Research & Training Institute (WRTI), Kenya for granting permissions to conduct the research. We are incredibly grateful to all the staff from WRTI from Tsavo East, Kenya Wildlife Service rangers, and Tsavo Trust staff and personnel for assisting with fieldwork and data collection.

## Ethical standards

This research abided by the *Oryx* guidelines on ethical standards. Fieldwork was conducted as part of regular monitoring by Wildlife Research & Training Institute and the Kenya Wildlife Service. The data involved no experimentation, collection, or monitoring of live animals.

## Data availability

Scat analysis data used in the manuscript can be found in Table 1. Scat locations, which also correspond to lion locations have not been added to the table because of their sensitive nature, and can be made available on reasonable request to the corresponding author. Prey abundance estimates are from published report.

## III. Figures and Figure Captions in the Main Text & Supplementary Information

**Figure S1.**
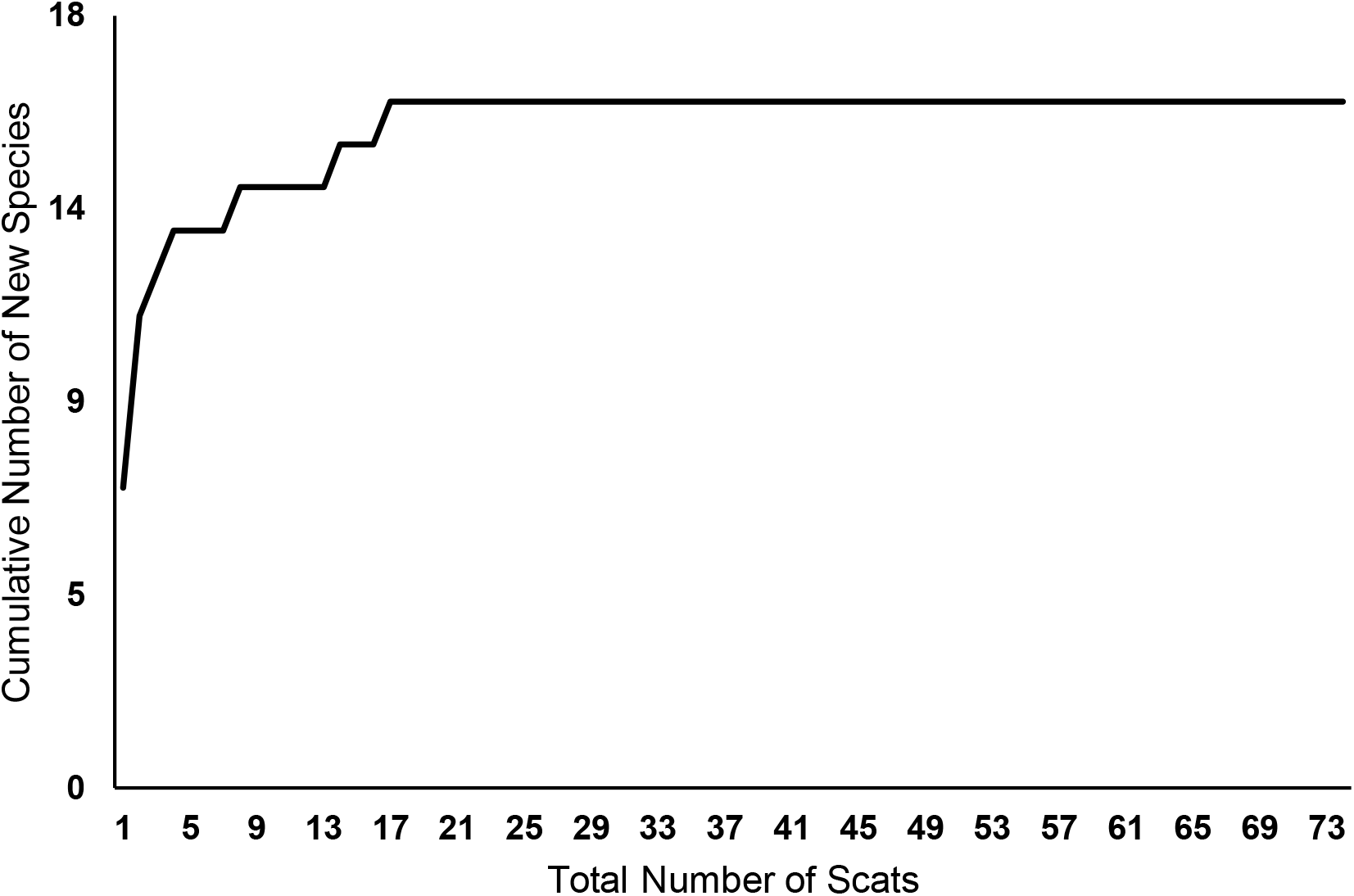
Calculation of sampling adequacy of lion scats from Tsavo East National Park, Kenya 2019-2023 for the characterization of diet, with new prey satiating at ∼20 scats.

